# Prolonged body temperature tuning in mice using radio frequency generating electromagnetic resonant circuit-system

**DOI:** 10.1101/2023.04.30.538865

**Authors:** Grégory Franck, Marie Le Borgne, Hélène Cazier, Patrick Laignel, Giuseppina Caligiuri, Antonino Nicoletti

## Abstract

**Rational:** In endothermic animals, body temperature (BT) is an evolutionary conserved and well characterized physical parameter that guarantees physiological functioning state. It results from the sum of bioenergetic processes of the body weighted by behavioral strategies, heat loss, and thermolytic processes. However, the intrinsic impact of temperature and temperature changes on the biology is far less understood. To date, the modification of the environmental temperature has constituted the main lever to evaluate the impact of thermic changes in small animal. However, studying intrinsic effect of temperature remains impossible using conventional laboratory equipment, mainly because hypothalamus instructed with information grabbed from the environmental temperature finely regulates and maintain body temperature around 37°C. Numerous pharmacological treatments have been used to block these thermoregulatory mechanisms, but confer high toxicity while dysregulating the central nervous responses and can potentially have confounding direct effects on studied peripheral tissues. Alternatively, physical methods using energy irradiation were reported, but they remain expensive and usually involve animal immobilization. We aimed at designing a simple and affordable device to adjust and maintain body temperature on the long course in conscious and free-moving animals.

**Method:** We developed an electromagnetic LC resonant circuit (ELM circuit) producing a radio frequency signal (64 kHz) inside a copper coil refrigerated with a water circuit. This setting is powered by a simple a 0-48V AC generator, allowing the use of a domestic electrical network. This setting can accommodate metal-free 3D-printed circular cages, where adult mice, previously implanted with thermometric ID transponders, are monitored remotely for intraperitoneal temperature over time.

**Results:** The BT of mice placed in the ELM circuit could be regulated in a reproducible fashion. Healthy mice increased their BT from 37 to 39.8±1°C, upon power supply ranging from 0 to 48V, respectively. In septic mice developing hypothermia (33±1 °C), BT could be either normalized to normothermia (37°C, 24V), or increased to fever-range hyperthermia (40°C, 48V) as a function of radiofrequency energy. BT tuning was accurate and stable for at least 12h. Blood count after 6 or 12 hours showed no modifications between groups, cardiomyocyte displayed heat shock response within the first hour in mice exposed to the maximal dose (BT=41°C). MALDI TOF imaging on brain microsections revealed modifications of the brain proteome, as suggested by differential PKC-theta, and prolactin 7B1 load in heated mice, as compared to controls.

**Conclusion:** Precise body temperature tuning is achievable in small animals, and could be of high interest to understand the impact of temperature in (patho)physiology.

## Introduction

Temperature in the Universe ranges from -273°C up to 1 000 000°C. With few exceptions, living organisms can only withstand a very narrow temperature range between 0 and 45°C. Life is therefore intrinsically dependent on temperature. This is primarily due to the state of water from which biological reactions rely. In mammals, most of which are endotherms, body temperature (BT) is very tightly regulated within a narrow range because most, if not all, biological activities are temperature-dependent. Any deviation from this range alters the normal functioning of cells and tissues. This is particularly the case during inflammatory responses, which are most often associated with hyperthermic responses (either locally or systemically with fever), but can also cause paradoxical hypothermic responses.

Fever is a common manifestation in endotherm animals and is considered a protective mechanism in the case of infection, where an increase of BT of 1°C to 4°C appears to improve clinical outcome. It is interesting to note that both ectotherms and plants have adopted similar strategies (but through distinct mechanisms) to raise their core temperature to face infectious agents. A classic explanation is that febrile temperatures reduce the replication potential of pathogens and/or make it easier to lyse them. There is also now a growing body of evidence indicating that fever-like temperature can boost the effectiveness of the immune response by acting on both innate and adaptive immune effectors (see ^1^ for review).

However, fever is not always beneficial. In situations of intense inflammation, lowering rather than raising BT can be a protective strategy. This is seen in patients with sepsis or neurological injuries, where uncontrolled fever is associated with worse outcomes, while treatments that induce hypothermia have been shown to have clinical benefits ^2,3^.

A challenge in evaluating the precise impact of BT in endotherms comes from the drawbacks of existing experimental models. The first approach used in many experimental studies examining the impact of temperature *in vivo* is to manipulate the environmental temperature using heating pads, cooling pads, temperature-controlled rooms, or baths. These studies have found deep modifications of the metabolic needs of mice. For example, thermoneutrality (typically around 30-32°C), a condition in which mice spend the least amount of energy to adjust their BT, has been found to alter glucose metabolism, modulate immune functions, reduce lipid metabolism and potentiate atherosclerosis compared to ambient temperature conditions ^4-7^. Exposure to chronic cold stress also promotes the alteration of innate immune functions ^8^, and the growth of atheromatous plaque ^9^. In both opposite experimental conditions, the hypothalamus, instructed with information on environmental conditions, quickly and accurately regulates BT in response to changes in external temperature.

Pharmacological methods have also been reported to either increase (adrenergic and cholinergic agonists, TRPV1 agonists, prostaglandins, catecholamines, cannabinoid, and thyroid hormone) or decrease BT (adrenergic or cholinergic antagonists, TRPA1 agonists, non-steroidal anti-inflammatory drugs, barbiturates, benzodiazepines, and opioids). Pyrogenic endotoxins such as LPS can cause hypothermia or fever, depending on their dose of administration and on ambient temperature ^10,11^. However, these methods introduce a plethora of confounding mechanisms, such as aspects of the inflammatory response, which prevent a precise understanding of the impact of temperature *in vivo*.

Other methods to maintain elevated temperatures have been reported, including the ectopic application of laser beam or infrared energy ^12,13^. These approaches are complex to implement and require in most cases the animal to be anesthetized or restrained. While anesthesia generally causes a decrease in BT, restraint generates excessive stress and rather leads, in the absence of habituation, to a significant increase in BT. Alternatively, microwave irradiation was reported to induce transient whole-body hyperthermia, but required the mice to be anesthetized ^14^.

Studies tackling the intrinsic impact of temperature have thus shifted to controlled protocols *in vitro* ^15^. However, this approach does not allow to study the impact of temperature in patho-physiologies involving a broad range of cellular actors. In this context, we aimed to develop a simple, safe and affordable device to investigate the intrinsic effect of temperature in mice during prolonged protocols.

## Material and methods

### Water-cooled LC resonant circuit

A 6 mm diameter T2 copper tubing was bent to form a 21 cm diameter coil composed of 3 spires. The LC resonant circuit was built with a L value = 2.49 μH (coil inductance), and a capacitance C = 1.98 µF (6 * 0.33μF capacitors in parallel, MKPH 1200v 50Kz), to deliver a 64 kHz induction field. Components were mounted on a zero volt switching (ZVS) type circuit, to generate a high intensity and high frequency current from a direct current source. It includes MosFet type transistors (IRFP260N.) whose gate voltage is limited by a 12v Zener diode (5A), allowing a variable voltage supply up to 48v to vary the intensity of the magnetic field. This ZVS circuit induces an eddy current in the mass of an object placed in the coil, causing a voltage-dependent heating of the object. A water pump equipped with a cooling thermostat (Lauda, RE 630 SN) was connected to the copper coil inlet and outlet, using a 10 mm silicon tubing (Avantor), allowing the coil to be cooled by water circulation. All the electronic boxes were printed in 3D stereolithography with an anti-electrostatic resin (ESD Resin, Formlabs), and cages were printed with transparent resin (Clear V4, Formlabs), using the Form 3L SLA printer (Formlabs). This system the invention is protected by registered patent referenced as Franck G, Le Borgne M, Laignel P, Nicoletti A, “System and method for controlling the body temperature of an animal in a cage” BET 23L0989/ECE, April 2023.

### Mice

Eight to twenty-week old C57BL/6N mice (Janvier Labs) were fed a regular chow diet. All investigations on mice conformed to the Directive 2010/63/EU of the European Parliament, and review and approval of the study was obtained from the *Comité d’Ethique Paris Nord #121* (APAFIS #9687, #18976, #30125, and #34536).

### RFID and iron pad implantation

Mice were anesthetized with isoflurane, and implanted intraperitoneally with sterilized RFID transponders (TAM-L/temperature, ISO FDX-B), measuring 2.12 × 13 mm, allowing instant and wireless animal identification, as well as fast and noninvasive temperature measurement. For some experiments, irons pads (1cm x 1 cm) were surgically implanted concomitantly with FRID transponder implantation. Muscle and skin closure was then performed using surgical sutures. Mice were then allowed to rest for a week before the start of any experiments.

### Mouse RF exposure

Up to 5 previously habituated mice were housed in each cage, with food and water ad lib. A 64 kHz radiofrequency exposure was applied, and mice were monitored for up to 12 hours. Wireless thermometry using implanted transponders was performed every 15 to 30 minutes. Cage and room temperatures were monitored continuously. At the end of the experiment, retroorbital blood sampling was performed on EDTA, and mice were killed by isoflurane overdose. Compete blood count was measured on Vet ABC (Scil).

### Cecal slurry preparation and injection

Sepsis in mice was induced by injecting (i.p.) 2mg/g cecal slurry. Cecal slurry was prepared as previously described ^16^. Briefly, the whole cecum content from 10-15-week-old C57Bl/6 mice was extracted, weighted, mixed with sterile water at a ratio of 0.5-ml of water to100-mg of cecal content, and filtered through sterile meshes (70µm). The resulting mixture was then mixed with an equal volume of 30% glycerol (in PBS), stirred, dispensed into cryovials, slowly frozen down to -80°C, and conserved up to 6 months. Following its thawing, cecal slurry then injected in mice, at a rate of 2mg/g of mouse.

### IR-thermometry

Mice implanted with an iron pad were imaged with an infrared camera (FLIR A300). For in vivo experiments, mice were anesthetized using xylazine/ketamine (i.p.) and the skin of the belly was shaved. Mice were then placed on their backs and imaged for a few minutes.

### Flow cytometry analysis

Spleens and bone marrows from tibias were dissociated in PBS on a 100-µm cell strainer, then incubated in Ammonium-Chloride-Potassium lysis buffer for 5 minutes at room temperature to lyse red blood cells. Cells were counted with a Scepter Cell Counter. Cells were incubated with a purified rat anti-mouse CD16/32 (clone 2.4G2, Fc Block, BD Biosciences) for 10 minutes at 4°C. Cells were then incubated for 20 minutes with a mix of antibodies (Table 1). Data was acquired on a LSRFortessa X-20 flow cytometer (BD Biosciences) and analyzed using FACSDiva (BD Biosciences) and FlowJo (TreeStar) softwares.

**Table 1:** Antibodies for flow cytometry.

| Target | Fluorochrome | Isotype | Clone | Provider | Reference | RRID |
| --- | --- | --- | --- | --- | --- | --- |
| Mouse CD3 | BV421 | Rat IgG2b, κ | 17A2 | BD Biosciences | 564008 | AB_2732058 |
| Mouse CD4 | PE | Rat IgG2b, κ | GK1.5 | BD Biosciences | 557308 | AB_396634 |
| Mouse CD11b | BV711 | Rat IgG2b, κ | M1/70 | Biolegend | 101241 | AB_11218791 |
| Mouse CD19 | BUV737 | Rat IgG2a, κ | 1D3 | BD Biosciences | 612791 | AB_2870111 |
| Mouse CD45 | BV605 | Rat IgG2b, κ | 30-F11 | BD Biosciences | 563053 | AB_2737976 |
| Mouse CD193 | PE-Cy7 | Rat IgG2a, κ | J073E5 | Biolegend | 144514 | AB_2565740 |
| Mouse F4/80 | A488 | Rat IgG2a, κ | BM8 | Biolegend | 123120 | AB_893479 |
| Mouse Ly-6C | APC-Cy7 | Rat IgG2c, κ | HK1.4 | Biolegend | 128026 | AB_10640120 |
| Mouse Ly-6G | APC | Rat IgG2a, κ | 1A8 | BD Biosciences | 560599 | AB_1727560 |

### Immunodetection of soluble molecules

Plasma was prepared from blood samples, aliquoted and frozen at -80°C until use. Soluble molecules were quantified using ProcartaPlex multiplex immunoassay kits (Invitrogen) according to manufacturer instructions. Beads were analysed on a Bioplex-200 analyser (Bio-Rad).

### Statistical analysis

Statistical analysis were performed using Prism software. Mann-Whitney tests were conducted to compare two populations (as n<20). Restricted maximum likelihood (REML) analysis was conducted to compare groups in kinetics experiment. P-values are indicated when p<0.01. Quantitative data are expressed as mean +/-standard error.

### MALDI Mass Spectrometry Imaging

MALDI - MSI (Matrix Assisted Laser Desorption Ionization – Mass Spectrometry Imaging) analysis were performed on 3 µm thick serial section mounted on ITO (Indium Tin Oxide) poly-L-Lysine coated conductive slides. Successive slices were deposited on Superfrost slides and stained with HES to correlate MALDI images with histological features. Slides were dried at 37°C overnight before analysis. For MSI analysis, slides were pre-heated at 85°C for 15 minutes and dewaxing was done with xylene (2×5 min). Slices were rehydrated with successive bath of isopropanol (100%) and ethanol (100, 96%, 70% and 50%) for 5 min each. Antigen retrieval was performed using a decloaking chamber (BioCare Medical, Concord CA) by heating the section in H_2_O at 110°C for 20 min. Trypsin solution (25µg/mL, Promega France), 20 mM ammonium bicarbonate and 0.01 % glycerol (Sigma Aldrich, St Quentin Fallavier, France) was deposited onto tissue sections using an automatic sprayer (15 µL/min, 16 cycles, TM Sprayer, HTX Technologies). The incubation time for digestion was 3 h at 50°C in a wet chamber with a saturated solution of K_2_SO_4_. CHCA (α-Cyano-4-hydroxycinnamic acid, Sigma Aldrich, St Quentin Fallavier, France) MALDI matrix (10 mg/mL in Acetonitrile/H2O, 70/30 v/v, 1% TFA (Trifloroacétique acid) application step was also done with TM sprayer (120µL/min, 4 cycles). MALDI-TOF acquisitions were performed in positive ionization with the reflectron mode using the Smartbeam laser of the Autoflex III mass spectrometer. Data acquisition were performed using FlexControl 3.4 and FlexImaging 4.1 software packages (Bruker Daltonics) in the range of m/z 600–3200 at spatial resolution of 50 μm. A peptide calibration standard mix including angiotensin II, angiotensin I, substance P, bombesin, ACTH clip 1-17, ACTH clip 18-39, and somatostatin 28 (Bruker Daltonik GmbH) was used for external calibration.

### MALDI Data processing

Data analysis of the MALDI imaging data was performed with SCiLS Lab Pro 2023 (Bremen, Germany). The raw images were loaded in the SCiLS Lab software and baseline subtraction was performed using the convolution algorithm and spectra were normalized by total ion current (TIC). The peaks were then aligned using specific tool provided by Bruker [15]. Segmentation was performed using the software pipeline with weak denoising for the brain slice area of the 37°C and 40°C exposed mice. Segmentation corresponds here to a hierarchical clustering which groups together same spectra meaning that areas with same peptide profiles, and so similar protein tissue composition, are represented under a same color. PCA (Principal Component Analysis) was achieved based on the segmentation pipeline peaks list, working on all individual spectra with weak denoising and unit variance scaling. Based on the same peptides peak list, proteins annotation was assessed using the online version of Mascot with 1 missed cleavage allowed.

## Results

### Technical parameters of the device

Inductance coils (**Fig. 1a and 1b**), setting (**Fig. 1c**), cages (**Fig. 1d**) and thermostatically controlled enclosure (**Fig. 1e**) were designed to allow reproducible and stable environmental conditions, as well as stable body temperatures. By using a water-cooling system, as well as a temperature- and humidity-controlled enclosure, the ambient temperature (AT) was not affected along the experiment and remained constant (AT = 20°C), despite the heating of the coils and electrical components (data not shown). Coil parameters shown in **Fig. 1a**, such as coil diameter (D = 210mm), height (h = 36 mm), tubing diameter (d = 6 mm), and number of spires (n = 3) define an overall inductance of the coil (L = 3.13 µH). Based on the overall capacitance (C = 1.98 µF), and the formula:

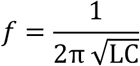

the theoretical frequency of the radio signal is *f* = 64 kHz (**Fig. 1b**). Two settings, arranged in series for the water-cooling system, were placed in a thermostated enclosure (**Fig. 1c**). Mouse cages, feeders, and bottle bases (**Fig. 1d**) were designed to avoid any magnetic interference, and thus avoid the detrimental increase of environmental temperature. A thermostated air flow was designed to limit ambient temperature derivation and temperature heterogeneity within cages. Calibrated iron pads (1 cm^2^) produced by 3D printed sintering aluminum (ProtoLabs) were used to evaluate the efficiency of the system. We observed a gradual heating of the pad as a function of the input voltage, with highest values becoming steady at +19°C (**Fig. 1f**). To assess potential biases caused by the localization inside the coil, we analyzed the effect of a displacement on the XY axis, and on the Z axis. While XY localization had low to no effect on the maximal temperature value of the pad subjected to a 12V pulse (**Fig. 1g**), Z axis strikingly affected steady-state temperature, with maximum values observed for pads at the center of the coil (**Fig. 1h**).

**Figure 1:**
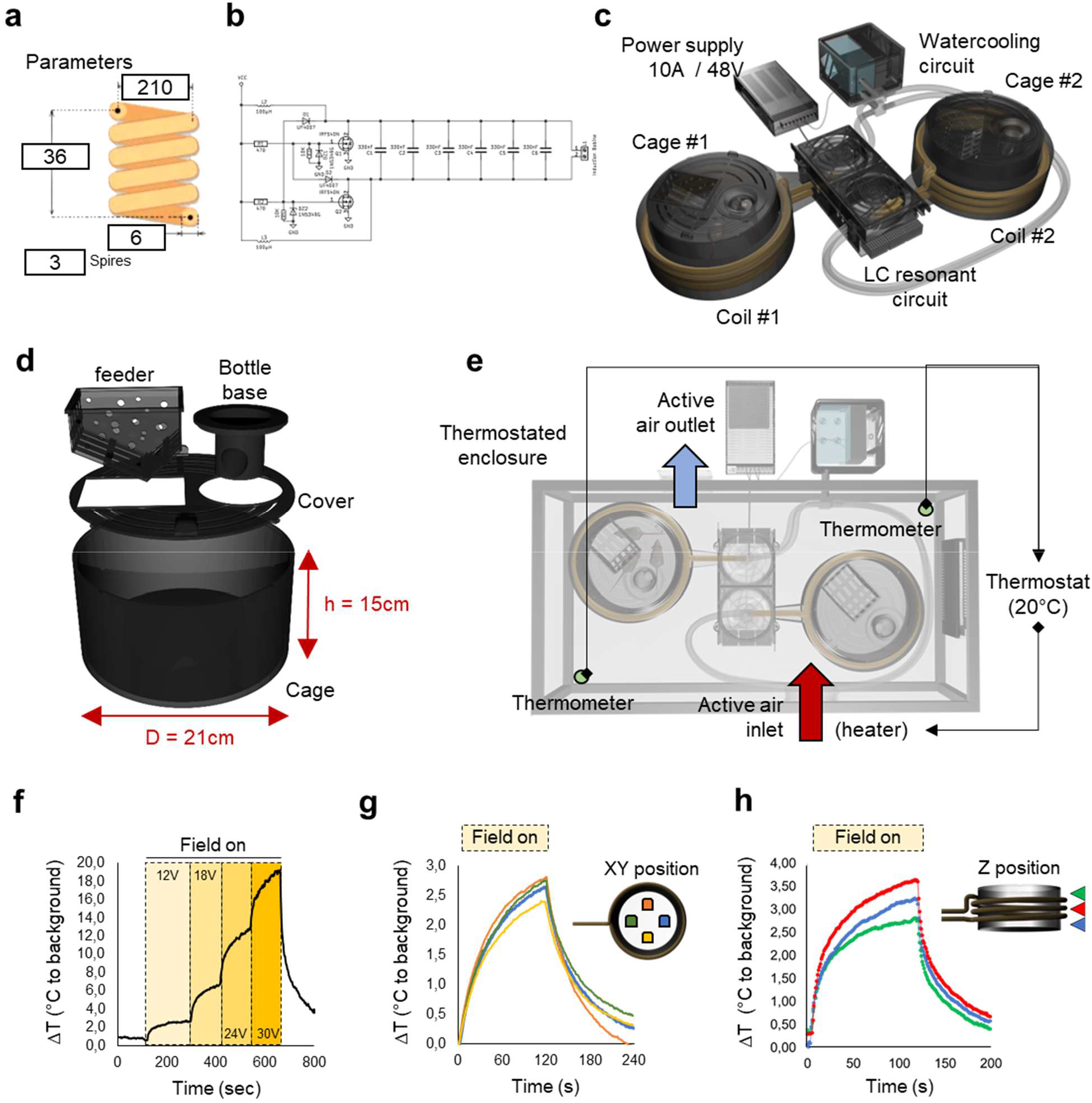
Electronic and cage design. **a**. Copper coil features aiming at generating specific impedance (µH) and signal frequency (Hz). **b**. Design of the LC circuit. VCC: voltage at the common collector; GND: signal ground; L: inductors; R; resistors; D: diode; Q: transistors; DZ: LEDs. **c**. Prototype schematic representation of two independent induction systems, serially connected to a water-cooling circuit. **d**. Dimensions of the compatible metal-free SLA-3D-printed cage system. **e**: Thermostated enclosure designed to control ambient temperature over experiments and time. **f-h**. Iron pad delta temperature (Δ°C, compared to non-ferrous background) assessed by IR-thermography according to input voltage (**f**), XY location (**g**) or Z localization (**h**) over time.

### Effect of RF exposure on mouse body temperature in physiological and pathological conditions

Mice previously implanted with RFID transponders (i.p.) were placed in the induction setting, and probed for internal BT for 6 to 12 hours (**Fig. 2a**). Short RF pulses with gradually increasing power resulted in increasing BT (**Fig. 2b**) as a non-linear function of the voltage imposed at the device terminals (**Fig. 2c**). Mouse BT rapidly decreased to baseline levels after RF shut down. Mice exposed to prolonged RF (48V) harbored sustained hyperthermia increase over 6 hours (**Fig. 2d and 2e**, 39.8±1°C vs 37°C in RF exposed vs control mice, respectively). BT increase was not correlated with body weight (**Fig. 2f**). To assess the possibility of modifying BT in septic animals, we injected mice with cecal slurry (2mg/g, **Fig. 2g**), leading to rapid and multiphasic hypothermia (**Fig. 2h**), accompanying animals until death (data not shown). Mild RF exposure (24V) allowed to adjust and sustain BT around basal temperature (37°C), while harsh RF exposure (48V) increased BT up to a febrile range (40°C) over the experiment (**Fig. 2h**). To induce local temperature increase *in vivo*, mice were implanted subcutaneously with aluminum pads. Maximal RF exposure (48V) induced an increase in their surface temperature above the pad implant, as assessed by IR thermography (**Fig. 2j**). Precise local temperature was then evaluated by implanting i.p. iron pads with a RFID transponder (**Fig. 2l**). While control mice reached a local temperature of 40°C during a maximal RF exposure (48V), mice with iron pads harbored a maximal local temperature of 42.5°C (**Fig.2m**). Altogether, these data show that the induction setting can increase BT in a precise and sustained fashion, in both normal or sick mice developing hypothermia, and that it can create a local hyperthermia through the use of metallic pads.

**Figure 2:**
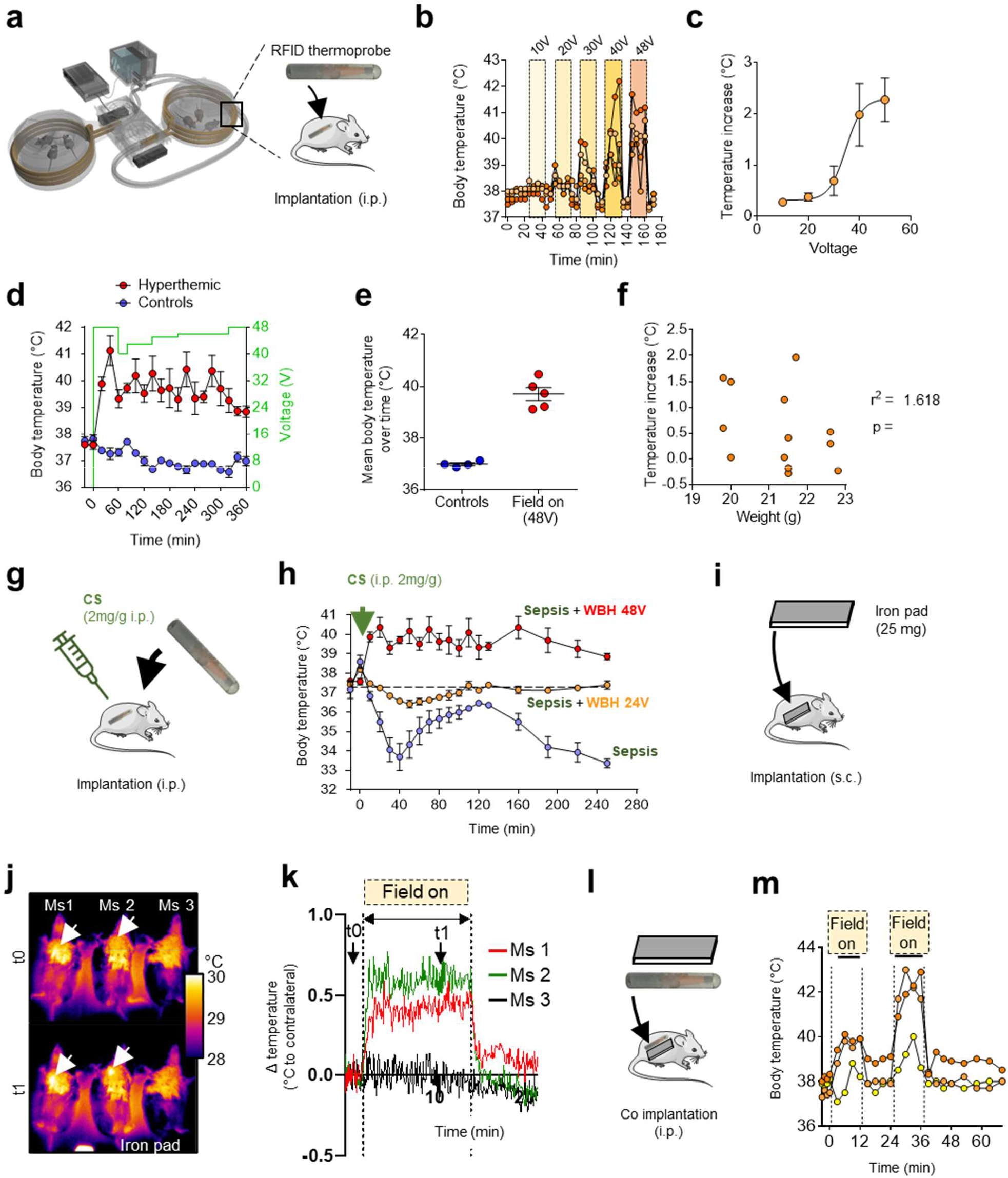
Control of body hyperthermia in healthy and septic mice. **a**. Experimental setting aiming at heating mice implanted (i.p.) with a RFID transponder before the experimentation. **b**. Effect of sequential and gradually increasing radiofrequency (RF) pulse on mouse body temperature (BT). **c**. Non-linear relationship linking voltage and BT. **d**. Body temperature in mice subjected to a 48V (RF) application for 6 hours (n=5) as compared to non-stimulated mice (n=4), and corresponding mean body temperature over 6 hours (**e**). **f**. Comparison between body temperature and body weight. **g**. Experimental setting with septic mice previously injected with a lethal dose of autologous cecal slurry (i.p). **h**. Body temperature of septic mice subjected to no RF, 24V, or 48V RF for 4 hours. **i**. Experimental setting aiming at heating mice implanted subcutaneously (s.c) with an iron pad. **j**. Infrared thermometry (IR) thermometry of anesthetized mice implanted (s.c) with an iron pad and subjected to maximal RD exposure (48V). **k**. IR skin thermometry over time and RF exposure. **l**. Experimental setting aiming at heating mice implanted (i.p.) with both an iron pad and a RFID transponder. **m**. Body temperature measured by the RFID probe during two RF pulses is shown for each mouse.

### Early heat shock response in cardiomyocytes after RF exposure

Temperature rise induces early heat shock in cells, characterized by the early translocation of heat shock factor 1 (HSF1) from the cytosol to the nucleus, to bind the heat shock response elements in DNA and promote the expression of HS related genes, including heat shock proteins. This response was transiently observed in the myocardium of mice exposed to RF, after 40, 50 and 60 minutes of exposure, while returning to baseline and unexposed controls levels after 70 minutes (**Fig. 3a and 3b**). These data confirm that RF exposure induces an increase in mouse BT, that is sensed in profound organs.

**Figure 3:**
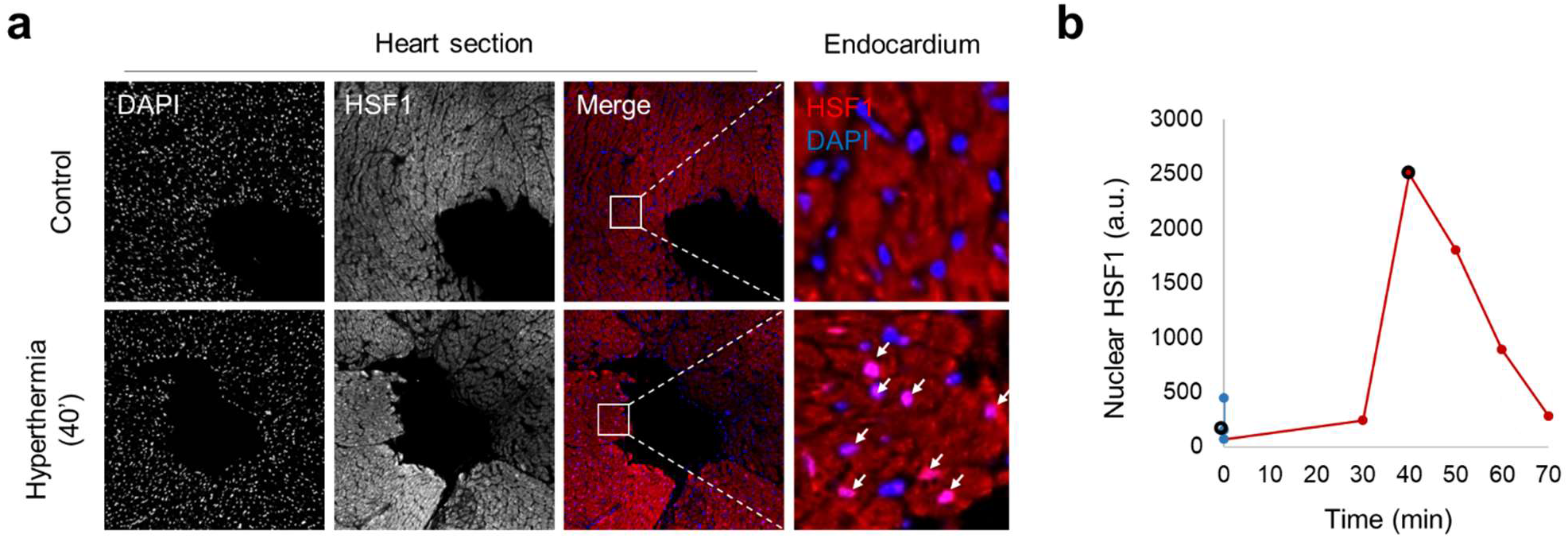
Effect of whole body hyperthemia (48V) on myocardial heat shock. **a**. HSF1 immunofluorescence (IF) in heart sections of mice subjected to a 48V radiofrequency (RF) exposure for 40 minutes (hyperthermia), as compared to control mice. **b**: Semi-quantitative nuclear HSF1 fluorescence in mice subjected to RF exposure during 30, 40, 50, 60, or 70 minutes (red dots), or in control mice (n=3, blue dots). Surrounded points corresponds to the IF images.

### Effect of RF exposure on blood count and inflammation

We investigated if RF exposure had a major impact on mouse health. Blood count and parameters were not affected by BT elevation, regardless of RF exposure time, even if a slight but not significant increase in red blood cell count and hematocrit was observed in mice exposed to RF for 48 hours (**Fig. 4a-c**). Mouse body weight was not affected during a 48h RF exposure, and mice did not develop any other clinical signs of pain during the exposure or in the week after (**Fig. 4d** and data not shown). Circulating IL-6 levels were found increased after a 6-hr maximal exposure (**Fig. 4e**), but showed no difference when mice were exposed for 48h, suggesting a transient elevation. Finally, 48-hr RF exposure did not modify the number of cells in secondary lymphoid organs (spleen and lymph nodes, data not shown), and the proportion of lymphoid and myeloid cells remained normal both in secondary lymphoid organs and in the BM (**Fig. 4f and 4g**). Therefore, apart from an early elevation of IL-6, these data demonstrate the absence of a significant impact of sustained hyperthermia on blood and inflammatory parameters.

**Figure 4:**
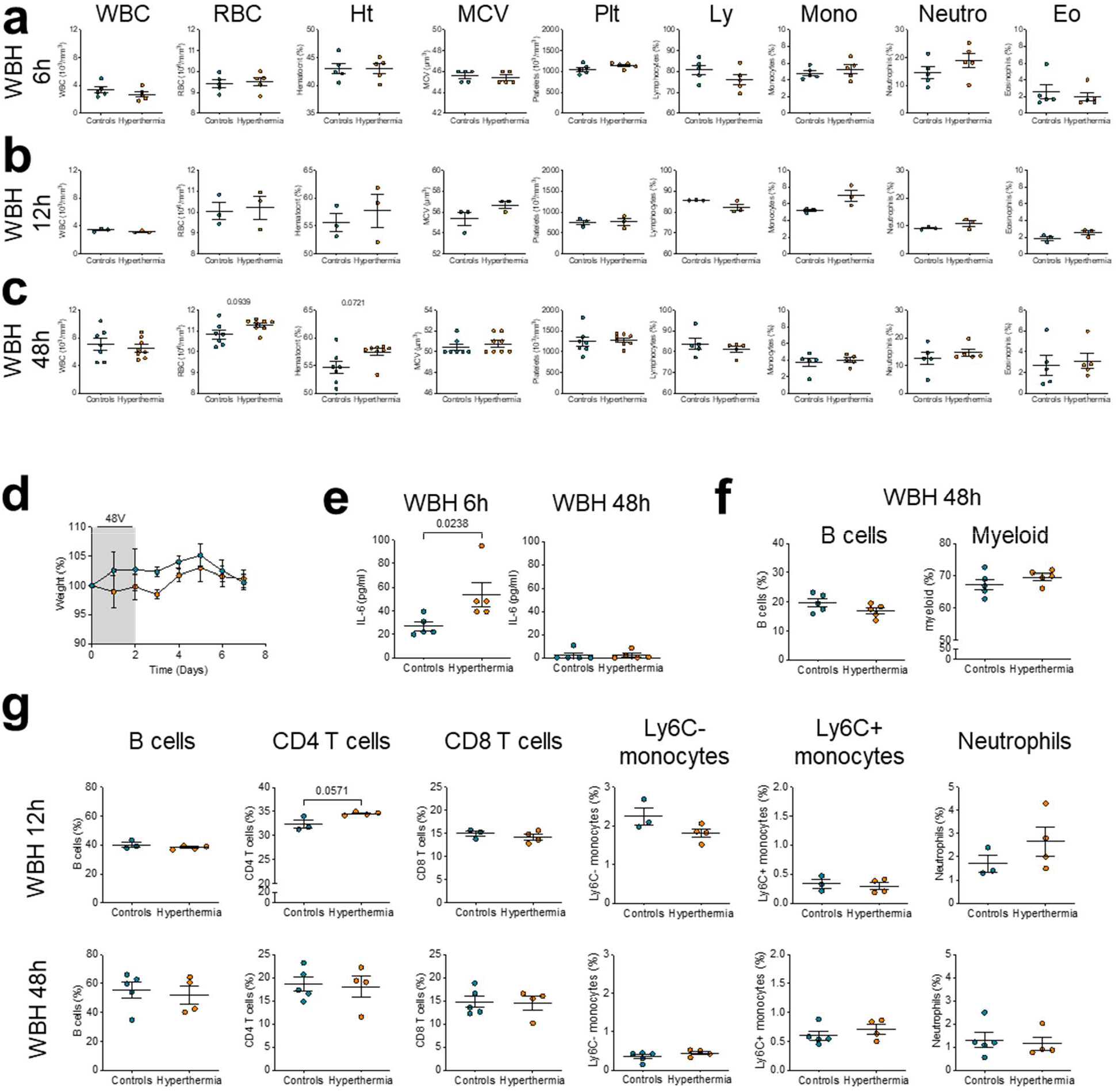
Effect of experimental body hyperthermia on blood count and systemic inflammation. Blood cell numeration performed on mice subjected to a 64 kHz radiofrequency exposure (orange dots) or control mice (blue dots) for 6 (**a**) 12 hours (**b**), or 48 hours (**c**). **d**. Body weight (no statistical effect of weight, REML analysis; n = 7-8 per group). **e**. Plasma IL-6. **f**. Spleen immune cell subsets (% in singlet CD45+ cells). **g**. Bone marrow immune cell subsets (% in singlet CD45+ cells). p-values of non-parametric Mann-Whitney test are indicated when <0.01.

### Spatial proteomic study in mouse brain in response to RF exposure

The brain constitutes the central regulator in response to thermal changes. Therefore, MALDI TOF imaging was carried out on this organ to evaluate the effect of whole-body hyperthermia (**Fig. 5**). Acquisitions were performed on brain coronal sections (Bregma: 0 mm) (**Fig. 5a**) from 3 mice exposed to RF exposure (average BT = 40°C) and 3 controls mice (average BT = 37°C). After the MSI processing of those whole-brain data acquired at 50 µm, segmentation was achieved to perform protein comparison. Mice exposed to a 6 hour RF exposure showed differential peptide distribution, as suggested by both principal component and ROC analysis (**Fig. 5b**). PCA was performed onto the white matter areas selected by segmentation in order to confirm the role of WBH on protein expression (**Fig. 5b**). A MASCOT search was performed using the peptide peak list from the segmentation results and showed that the putative regulated proteins with a fold change > 2 are involved in signaling, DNA repair, metabolism, trafficking, proteasome function and protein synthesis or endoplasmic reticulum stress (**Fig. 5c**). As an illustration, we highlighted 3 proteins of interest that were differentially expressed between RF-exposed and control animals: KIBRA (600.383 m/z ± 0.156 Da), PKC-theta (629.398 m/z ± 0.156 Da), and Prolactin (PRL) 7B1 (603.335 m/z ± 0.156 Da) (**Fig. 5d**). Immunofluorescent staining for PRL in coronal brain sections confirmed PRL overexpression in mice exposed to RF (**Fig. 5e**). Therefore, BT increase induced by RF exposure induces a major response in the brain.

**Figure 5:**
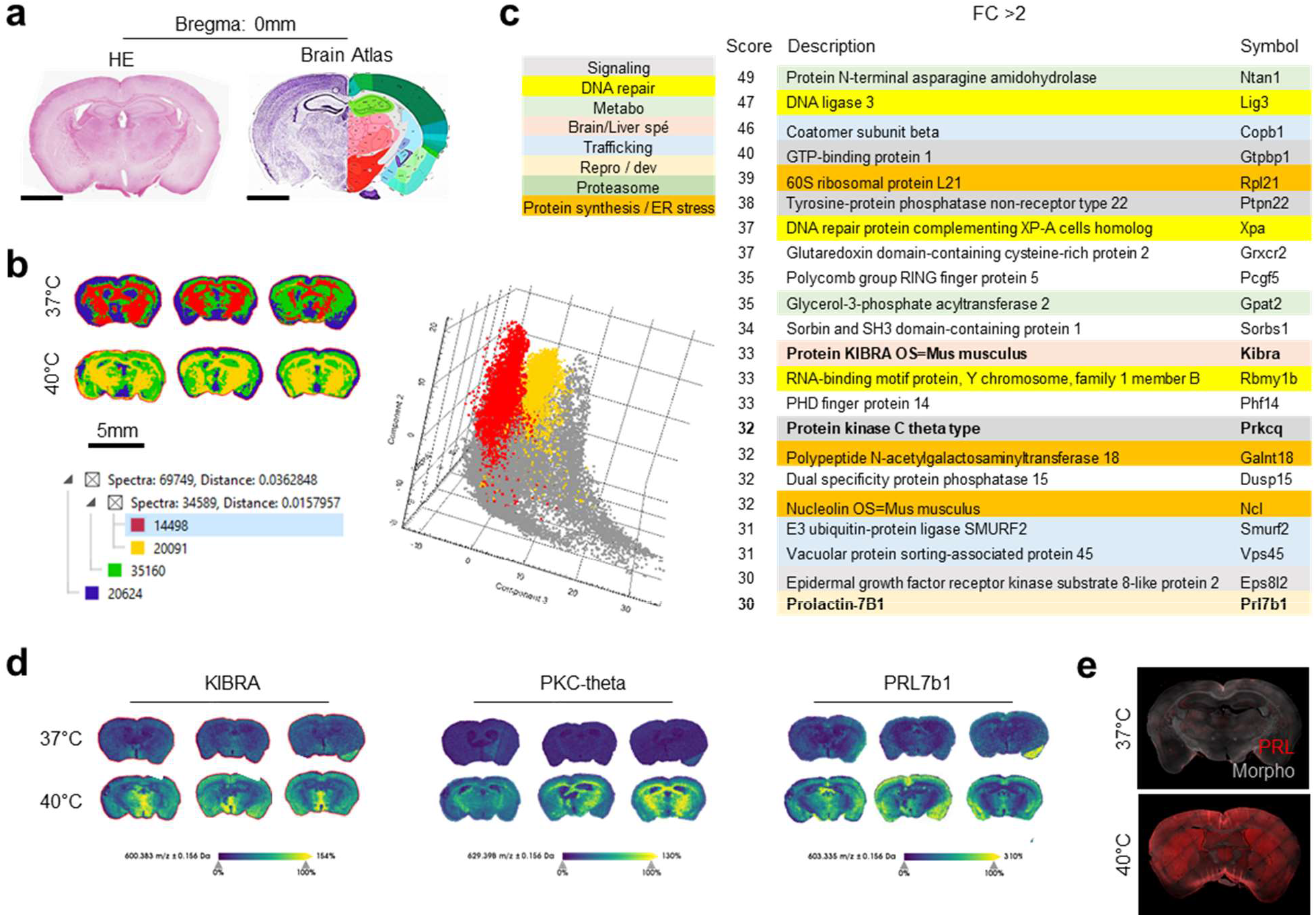
Effect of whole-body hyperthermia on protein expression profile obtained by spatial mass spectrometry. Matrix assisted laser desorption ionization (MALDI) coupled to time-of-flight mass spectrometry (MALDI-TOF MS) in coronal brain sections of mice subjected to radiofrequency (48V) exposure for 6 hours (group 40°C), or control mice (37°C). **a**. Coronal brain section plan used in the study with corresponding atlas (atlas.brain-map.org). **b**. Peptide differential expression assessed by principal component analysis (PCA) in segmented brain areas. **c**. Top 20 putative protein obtained by MASCOT with a fold change >2 between RF-exposed and control groups. **d**. m/z spectrum obtained for KIBRA, PKC-theta, and Prolactin-7B1 in brain sections of the two groups. **e**. Immunofluorescence of prolactin (PRL) in corresponding brains sections.

## Discussion

We introduce a new experimental device designed for studying the impact of temperature on small animals over extended periods. The device is simple to use, safe, and affordable, and employs induction heating. Induction heating, a method of generating heat through electromagnetic induction, is widely used in various industrial applications. In this process, an object is placed in a magnetic field that exerts a force on the free electrons present in the object, generating an electric current. The energy then dissipates inside the object in the form of heat. Our device adapts this method for controlling the BT of laboratory animals. This method has several advantages, such as the ability to rapidly control the temperature and the absence of physical contact, making it ideal for use in free-moving laboratory animal experiments.

BT results from a combination of factors, including the environmental temperature, as well as thermogenic and thermolytic processes, which can be driven either intrinsically or via behavioral modifications. When placed in a supra-thermic environment (above the thermoneutrality range), endotherms regulate their temperature through central mechanisms that control their physical activity and numerous involuntary responses that counteract temperature increase and lower BT. These responses include inhibition of adipose tissue thermogenesis, cutaneous vasodilation, sweating, and piloerection. They result from the integration of heat information sensed by nerve thermoreceptors in the skin that send a feedforward signal via the spinal cord and lateral parabrachial nucleus to the preoptic area (POA) in the hypothalamus (see ^17^ for review). Locally, the POA can also receive thermosensory information from hypothalamic neurons equipped with thermoreceptors, or pyrogenic signals such as prostaglandin E2 produced by the blood brain barrier and inflammatory cells in response to infection and inflammatory stimuli ^1^. Once integrated, these signals enhance the inhibition activity of the GABAergic projection neurons descending from the PAO. This leads to the inhibition of the motor adrenergic outputs, and results in cutaneous vasodilation and tissue thermogenesis inhibition (muscle and adipose tissue). Hence, studying the intrinsic impact of body temperature in animals by changing the environmental temperature is a true challenge since it associates with confounding and complex reactions. We believe that introducing an experimental setting that allows for changing the temperature of the tissues, but not of the environment, is an important milestone in this field.

We were able to show that with this experimental setting, the mouse BT can be increased by 3-5°C with very precise levels of the order of tenths of a degree by simply adjusting the potentiometer which controls the voltage supplied to the RF circuit. Moreover, this BT tuning could be achieved for prolonged period spanning several hours or days. Our design was optimized for mice but can be customized for smaller or bigger species. We analyzed whether the X-Y position of the animals in the cage could introduce variations and thus experimental biases. We found that X-Y have a minimal impact, with homogenous temperature elevation in iron pad exposed to RF. Instead, Z position significantly affected temperature changes, with the highest temperature recorded at the center of the coil. Therefore, cage localization on the Z axis should be carefully controlled between different cages and between sequential experiments, to ensure reproducibility.

Besides, in our setting, the animals are exposed to RF. The energy emitted during radiofrequency (RF) exposure is absorbed by the biological tissues, increasing the kinetic energy of the molecules, causing the molecules to vibrate more rapidly, triggering temperature elevation. The amount of RF energy absorbed by the tissue depends on several factors, including energy frequency, intensity, the duration of exposure, and the properties of the tissue itself. Some tissues, such as adipose tissue, absorb more RF energy than others, such as muscle. While our results did not show evidence that body weight influences temperature elevation during RF exposure, factors such as obesity, age and sex, that can affect the percentage of body fat, should be standardized in each experiment.

RF toxicity is still a matter of debates. While low level of RF exposure is not considered to be toxic, at higher levels of exposure, RF energy can cause thermal damage to the tissue, leading to burns and other types of injury. Additionally, long-term exposure to high levels of RF energy has been linked to an increased risk of cancer, particularly brain cancer ^18-20^. In clinic, high energy RF is used for therapeutic purposes, such as local hyperthermia for cancer treatment, where the objective is to damage or destroy the tumor cells. It was hence important to monitor several biological responses of the mice placed in our device for a prolonged period. First, we did not detect any macroscopic evidence of skin burns or organ damage. We found no effect on blood count or body weight besides a modest elevation of red blood cell count after a 48-hour exposure. At 6 hours, there was a transient and modest increase of circulating IL-6 that returned to basal levels at 48 hours. Other very slight and transient modifications in cell composition in the spleen and the bone marrow could be captured, that were found again in the normal range at 48 hours. We performed a deeper investigation on the effects that RF exposure could exert on the heart and the brain. When healthy mice were exposed to a maximal RF (48V), harboring a febrile range hyperthermia (40°C), we detected an early heat shock response in the cardiomyocytes. Again, this was a transient phenomenon, not observed anymore after 70 minutes of exposure. We also performed a comprehensive analysis of the proteome of the brain in mice exposed for 6 hours to a 48V RF circuit. The expression of several proteins was found to be altered, thereby indicating that the brain is extremely sensitive to temperature elevation. Yet, lipidomic profile should constitute of more relevant readout, since lipid mediators, such as prostaglandin E2, are key player in hypothalamic thermoregulation, as well as response to inflammatory and infection stimuli ^1^.

While fever is a common manifestation in endotherm animals, a paradoxical hypothermic responses can also be observed in some conditions. In humans, hypothermia is observed in 9 − 35 % of patients with sepsis ^21,22^ and has worse prognosis value than hyperthermia ^23^. With this, the underlying question of whether BT in critically ill patients should be managed arises. To evaluate if our device can tackle such questions, we tested its effect in septic mice mimicking this clinical situation. The physiological responses to infection can vary among different species, and the precise mechanisms underlying these responses are still not fully understood. During endotoxemia, mice develop hypothermia when housed at ambient temperature ^10^. This specific response could be explained by their high surface area to volume ratio, leading to fast heat loss compared to larger mammals. In patients, hypothermia could be explained both by microvascular dysfunction and vasoconstriction impairment, or by the inhibitory effect of PGE2 on heat production upon infection. In contrast, environmental temperature elevation has been found to induce fever in mice and rats (ref), but this response is not standardized between species. Here, the exposition to RF bypasses some of the neurophysiological regulations, since our device precisely tunes and maintains abnormal BT over time. Control mice that were placed in a zero V RF circuit had, as expected, an hypothermic response. Remarkably, we could impose a normothermic, or even a hyperthermic response thereby completely reverting their BT. This experimental setting is hence suited to study whether BT management can improve clinical outcome.

This device is also a promising tool to trigger local temperature elevation. Indeed, metal pads can be implanted to specific anatomical locations. As can be seen in our calibration experiments performed with metal pads, their heating is much more pronounced than the heating of mouse body; therefore lower voltage settings (20V), that only moderately increase the BT of normal mice, can heat iron pads by several degrees. It is hence possible to address the impact of temperature specifically elevated in a given organ, let it be the kidney or the liver for instance, or even lymph nodes.

